# Post-transcriptional control of SRSF9 promotes the epithelial-to-mesenchymal transition (EMT) in colorectal cancer cells

**DOI:** 10.1101/2022.05.16.492181

**Authors:** Chaitra Rao, Robert A. Svoboda, Siddesh Southekal, Heidi M. Vieira, Dianna H. Huisman, Deepan Chatterjee, Chittibabu Guda, Kurt W. Fisher, Olga A Anczuków, Robert E. Lewis

**Affiliations:** Eppley Institute, Fred & Pamela Buffett Cancer Center, University of Nebraska Medical Center, Omaha, NE 68198, USA; Department of Pathology and Microbiology, University of Nebraska Medical Center, Omaha, NE 68198, USA; Department of Genetics, Cell Biology and Anatomy, University of Nebraska Medical Center, Omaha, NE, University of Nebraska Medical Center, Omaha, NE 68198, USA; The Jackson Laboratory for Genomic Medicine, Farmington, CT, USA

## Abstract

In human colorectal cancer (CRC) cells the Raf/MEK/ERK scaffold Kinase Suppressor of Ras 1 (KSR1)-dependent signaling is required for the epithelial-to-mesenchymal transition (EMT)-like phenotype. Here we show that KSR1 promotes the association of differentially spliced mRNA bearing recognition sites for the Serine/Arginine-Rich (SR) splicing factor SRSF9. CRISPR/Cas9 disruption of KSR1 destabilizes SRSF9 protein, which interacts preferentially with mRNA encoding Epithelial Stromal Interaction 1 (*EPSTI1*). EPSTI1 protein mediates Ras and KSR1-dependent induction of EMT. Analysis of *EPSTI1* splice variants reveals that inclusion of exon 8 is critical to the ability of EPSTI1 to promote the E-to N-cadherin switch and CRC cell motile and invasive behavior. These data reveal a mechanism in CRC cells in which Ras-induced and KSR1-dependent signaling affects pre-mRNA splicing to control behaviors critical to cancer cell dissemination and metastasis.

## Introduction

The regulation of protein synthesis occurs at multiple levels including an intimate connection between transcription and RNA processing. All steps in RNA processing including RNA synthesis, pre-messenger RNA (pre-mRNA) splicing, intracellular transport, translation, and degradation, involve complex and regulated processes are mediated by RNA binding proteins (RBPs) that bind RNA to control its biogenesis and function [1]. pre-mRNA splicing is an essential process that involves removal of introns and inclusion or exclusion of exons from nascent pre-mRNA to produce mature RNA [2]. Unlike constitutive splicing, that involves removal of introns and ligation of exons in the sequential order they appear in a gene, alternative splicing (AS) allows a single gene to increase its coding capacity, allowing synthesis of structurally and functionally distinct protein isoforms. Each AS event is controlled by multiple RBPs, creating a cell-type specific distribution of alternatively spliced products. This dynamic RNA-protein complex has remarkable plasticity in substrate recognition and can influence many regulatory proteins bound to pre-mRNA to produce multiple mRNA isoforms [3, 4].

Splicing factors (SFs) are frequently altered in human diseases, leading to downstream changes in the spliced isoform repertoire [5–7]. Splicing factors are critical components of global gene expression and interact with unspliced and mature mRNAs to regulate all aspects of cell growth and development. Activator and repressor splicing factors bind specific sequences on their target pre-mRNAs eliciting concentration-dependent effects on splicing. Splicing factor binding is competitive, limiting the accessibility of another RBP that normally would repress or activate the splicing of that specific exon and thus elicit the opposite effect. Altered splicing-factor levels can expand or contract the exonic and intronic target sequences available, directly affecting the splicing of target exons by recruiting or repelling the splicing machinery. These observations have implications for normal cellular homeostasis and disease, including cancer. Transient changes in splicing-factor level, stability, or function can alter splicing to affect cell phenotype. [5, 7–12].

Cancer associated mis-splicing and post-transcriptional mechanisms reconfigure the proteome to control the cancer cell phenotype [6, 7, 12]. AS is a key step in post-transcriptional regulation that tightly balances epithelial and mesenchymal behavior during stem cell differentiation [12]. Several studies document the intricate connection between changes in AS and epithelial-to-mesenchymal transition (EMT) activation [11–14], highlighting how dysregulated AS could be a trigger for tumor relapse and metastatic spread, dramatically affecting patient outcome. Our current understanding likely reflects an initial understanding of the many mechanisms by which AS affects oncogenic signaling. Identifying splicing alterations in specific targets that contribute significantly to tumor formation and progression will facilitate the identification of new therapeutic targets.

SRSF9 is a member of Serine/Arginine-Rich (SR) proteins are a family of splicing factors (SRSF1 to 12) that act at multiple steps of spliceosome assembly, contributing to both constitutive and alternative splicing [7, 10, 15]. SR proteins have a modular structure consisting of one or two RNA-recognition motifs that determines their RNA-binding specificity, followed by a C-terminal domain rich in alternating serine and arginine residues (RS domain). Although SR proteins were initially described as activators promoting exon inclusion, transcriptomic studies suggest that they can promote inclusion for some targets and skipping for others [16]. SRSF9 displays a strong enrichment for the AGSAS motif (S= G or C) and is involved a wide range of functions, inclusion of SMN exon 7 [17, 18] and tau exon 10 [19], repressor of 3’ splice site selection of CE9 [20], exclusion of Caspase 2 exon 9 and CD44 exon 10 [21, 22], and 5’ splice-site utilization of Bcl-x_L_ [23]. SRSF9 was shown to promote ß-catenin protein synthesis and stimulate tumor growth *in vitro* and *in vivo* [24]. Although changes in several splicing factors have been shown to play a role in EMT-related process [12], how SRSF9 contributes to EMT during healthy development as well as in diseases remains to be determined.

In human colorectal cancer (CRC) cells the Raf/MEK/ERK scaffold Kinase Suppressor of Ras 1 (KSR1)-dependent signaling is required for the EMT-like phenotype [25]. We combined targeting of genes encoding key effectors of Ras signaling with polysome profiling to discover that KSR1 upregulates the translational efficiency (TE) of EMT-related transcripts. One of these targets, Epithelial Stromal Interaction 1 (*EPSTI1*), mediates Ras and KSR1-dependent induction of the switch in CRC cells from E-cadherin to N-cadherin [25], a central hallmark of EMT [26, 27] and anchorage-independent growth. In this study, we identified a novel KSR1-dependent signal promoting the stability of SRSF9. SRSF9 controls the E-to N-cadherin switch and is required for CRC cell migration and invasion *in vitro*. SRSF9 associates with *EPSTI1* mRNA and inclusion of exon 8 in EPSTI1 is optimal for induction of CRC cell EMT, suggesting that SRSF9-mediated splicing of *EPSTI1* is a Ras-induced and KSR1-mediated mechanism driving CRC motility and invasion.

## Results

### KSR1 and ERK activity are required for SRSF9 expression

KSR1-mediated signaling regulates the translational landscape of human colon tumor cells to support their survival [25, 28]. EPSTI1 and other mesenchymal mRNA translation are catalyzed by Ras-mediated KSR1-dependent ERK signaling. RNA binding proteins can direct mRNA transcripts onto polysomes and promote their translation to protein. To identify post-transcriptional regulators of KSR1-dependent translation control on EMT, we used the computational tool PRADA (<u>P</u>rioritization of <u>R</u>egulatory Pathways based on <u>A</u>nalysis of RNA <u>D</u>ynamic <u>A</u>lterations) [29] that identifies oncogenic RNA-binding proteins through the systematic detection of coordinated changes in their target RNA-binding motifs. PRADA uses gene expression representations of the RBP binding sites to scan the target sequences of interest for the presence or absence of matches. Since many RBPs fall into families with highly similar binding preferences, PRADA analysis introduced an additional penalty term that excludes RBPs whose activity is unchanged in the analysis, which extends and stabilizes the lasso regression model. We applied PRADA to our dataset to identify RBPs that had predicted binding sites to KSR1-dependent mRNAs. PRADA predicted binding sites for the top eight RBP affected by KSR1 knockdown (**Figure 1A**). Among the eight RBPs, the size and the direction of the regulatory potential assigned to consensus binding sites of SRSF9 were disproportionately decreased in the absence of KSR1. This analysis predicted only SRSF9 as a candidate RBP and no other SR family members such as SRSF1, SRSF2, SRSF7 and SRSF10. Analysis of SRSF9 expression in normal and tumor colon tissue from publicly available databases supports the role of SRSF9 contributing to tumorigenesis. The mRNA expression from TCGA and protein levels from CPTAC shows a significant upregulation of SRSF9 mRNA and protein colon adenocarcinoma compared to normal counterpart (**Figure 1B**). We tested the effect of KSR1 depletion on SRSF9 in CRC cell lines. SRSF9 protein expression was decreased upon KSR1 depletion in HCT116 and the expression of SRSF9 was restored in knockout cells upon expression of a KSR1 transgene (+KSR1) (**Figure 1C**). Treatment with ERK inhibitor SCH772984 [30] displayed a modest suppression of SRSF9 protein expression in tumorigenic patient-derived colon organoids engineered with deletion of APC, p53, SMAD4 and K-Ras^G12D^ mutation (K/A/P/S) (**Figure 1D**). These results suggest that ERK and KSR1 activity is crucial for SRSF9 expression in colon tumor cells.

**Figure 1.**
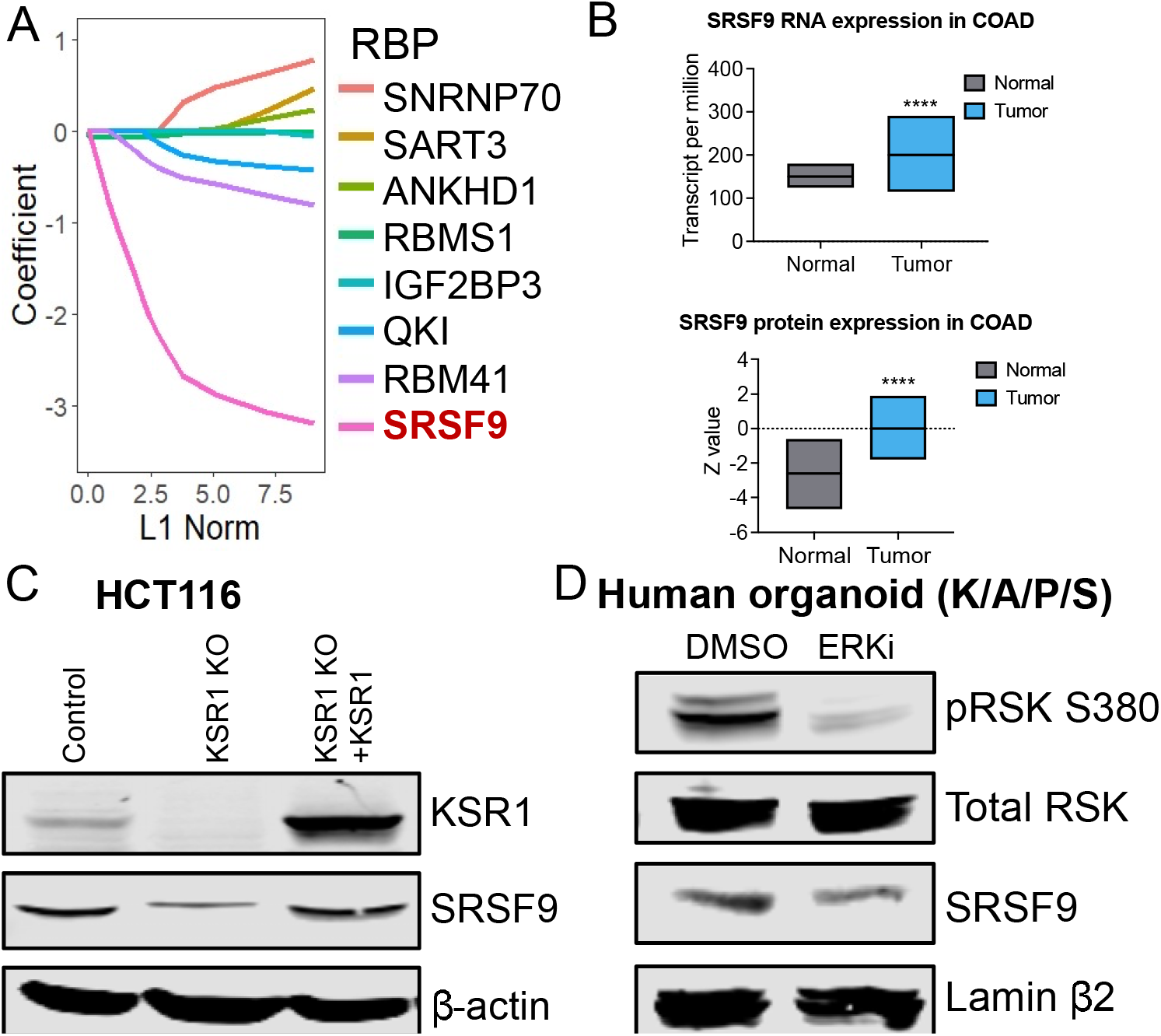
KSR1 and ERK activity are required for SRSF9 expression. (A) SRSF9 interaction with differentially translated mRNAs is suppressed by KSR1 RNAi. Regression coefficients of the indicated RNP as set by PRADA. Data reflect the change in coefficient caused KSR1 depletion. (B) RNA and protein expression of SRSF9 in normal and colon adenocarcinoma (COAD) from (top) TCGA and (bottom) CPTAC obtained by the UALCAN portal and plotted using GraphPad Prism. ****, P < 0.0001 (C) Western blot analysis of KSR1 and SRSF9 in HCT116 cells. (D) Colon tumoroids (K/A/P/S) ± ERK inhibitor SCH772984, were probed for phospho-Rsk (S380), Rsk, SRSF9, Lamin β2.

### KSR1 prevents SRSF9 protein degradation

To assess whether KSR1 regulates SRSF9 mRNA expression, HCT116 and SW480 control and KSR1 knockout (or HCT116 knockdown) cells were assessed for SRSF9 mRNA levels by RT-qPCR. Data form three biological replicates (each measured in triplicates) are shown. Depletion of KSR1 did not affect SRSF9 mRNA expression in both HCT116 and SW480 cells (**Figure 2A-B**). The RS domain of SR proteins are extensively phosphorylated on the serine residues, and it plays an important role in regulating subcellular localization and the activities of SR proteins. AMPK was shown to phosphorylate SRSF9 [31] and nuclear stress bodies repress splicing through CLK1-mediated phosphorylation of SRSF9 during heat stress recovery [32].. However, we detect no evidence of ERK-dependent SRSF9 phosphorylation (data not shown). SR proteins are also modified by the covalent attachment of polypeptides such as ubiquitin (Ub), and these Ub chains can target SR proteins for degradation by 26s proteasome. PhoshoSitePlus database [33, 34] identified Lys28 and Lys36 as two potential ubiquitination sites in SRSF9. On the basis of these observations, we tested whether depletion of KSR1 is promoting ubiquitin mediated SRSF9. We assessed SRSF9 expression following proteasome inhibition with MG132 for 6 hours. SRSF9 levels were rescued upon MG132 in KSR1 knockout cells indicating that KSR1 contributes to stabilization of SRSF9 (**Figure 2C**). We also examined SRSF9 turnover in HCT116 cells following treatment with cycloheximide (CHX) with and without KSR1 expression. Representative western blots of HCT116 cells from two independent experiments are shown (**Figure 2D**). Data from each experiment was quantified and the half-life of SRSF9 in each condition was calculated. Depletion of KSR1 reduced the rate of SRSF9 turnover. SRSF9 half-life was 5.5 hours in control cells but was reduced to 1.5 hour in KSR1 KO HCT116 cells. Together with these results suggest the KSR1 prevents the degradation of SRSF9 in colon cancer cell lines.

**Figure 2.**
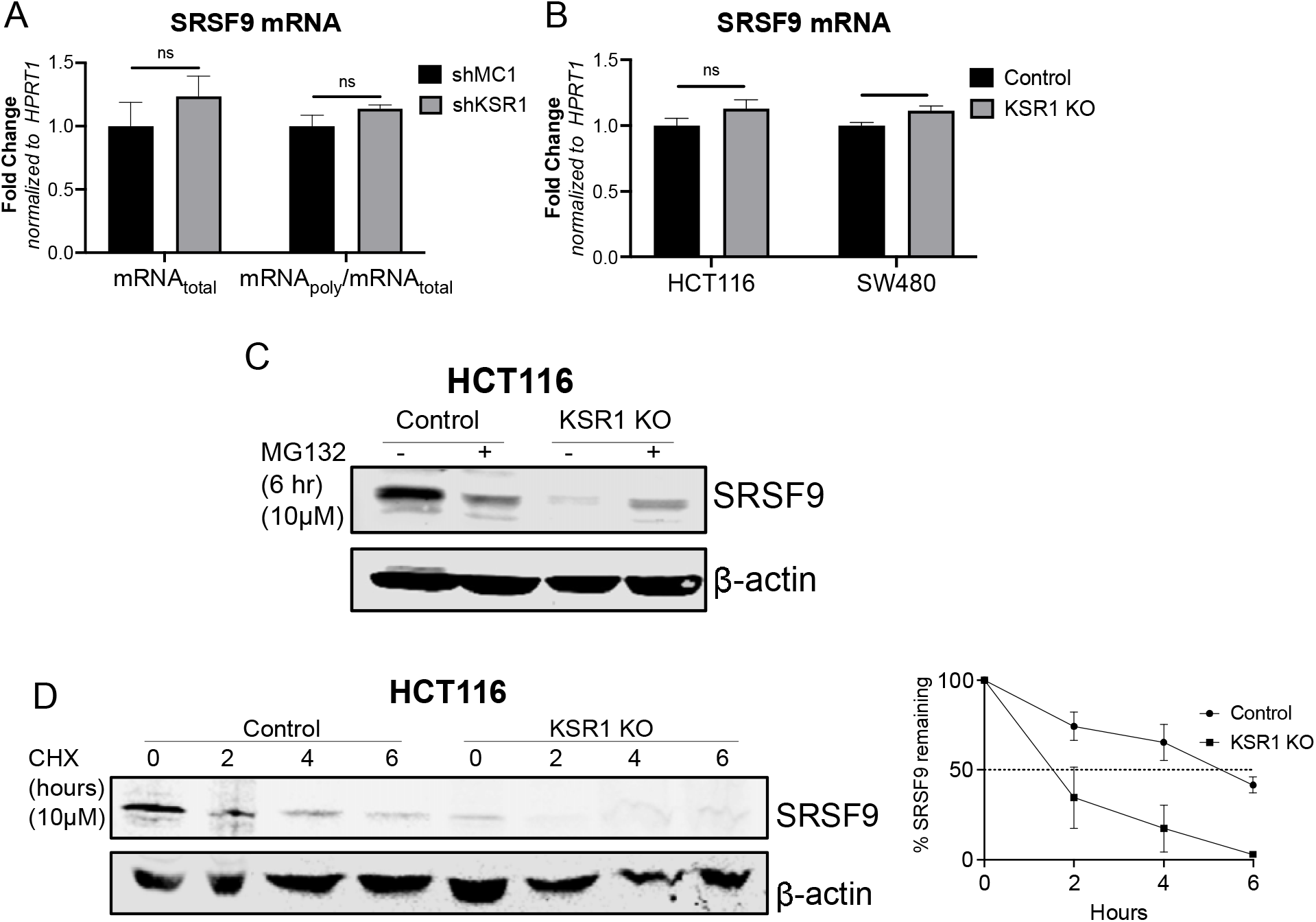
KSR1 inhibits SRSF9 protein degradation. (A) RT-qPCR analysis of SRSF9 mRNA from total RNA and polysomal RNA (fractions number 6-8) in control and KSR1 knockdown HCT116 cells, the TE was calculated as the ratio of polysomal mRNA to the total mRNA (n=3; ns, non-significant). (B) RT-qPCR analysis of SRSF9 mRNA from total RNA in control and KSR1 KO HCT116 cells (n=3; *, P<0.01; ns, non-significant) (C) Control and KSR1 KO HCT116 cells were treated with or without 10 μM proteosome inhibitor MG132 and blotted for SRSF9 and β-actin (D) Control and KSR1 KO HCT116 cells were treated with 10 μM cycloheximide for the indicated times. SRSR9 and β-actin were detected by western blot. (Bottom) SRSF9 expression at 0 hour was normalized to 100% and SRSF9/b-actin was quantified on a Li-Cor Odyssey.

### SRSF9 promotes CRC cell motility and invasion

SRSF9 was shown to promote cell invasion and migration in breast cancer models but does not impact cell proliferation or cell death [11]. The ability of KSR1 to regulate the migration and invasion in colon cancer cells, suggests that likely SRSF9 functions phenocopy that of KSR1. To test this possibility, we knocked down SRSF9 expression in HCT116 and SW480 cells using siRNA (**Figure 3C**) and subjected the cell monolayers to an *in vitro* scratch wound assay as previously described [25]. Targeting SRSF9 significantly impaired migration to repair the wound in monolayer by over 50% (**Figure 3A**). We assessed the role of SRSF9 contributing to invasive capacity in colon cancer cells, Transwell invasion assays through Matrigel in HCT116 and SW480 cells showed that SRSF9 disruption significantly suppressed invasion by over 75% (**Figure 3B**). Consistent with these data, SRSF9 RNAi inhibits N-cadherin expression with a coincident increase E-cadherin expression in HCT116 cells (**Figure 3D**). These data reveal a novel KSR-dependent signaling pathway regulating SRSF9 in EMT-like phenotype in colon cancer cells.

**Figure 3.**
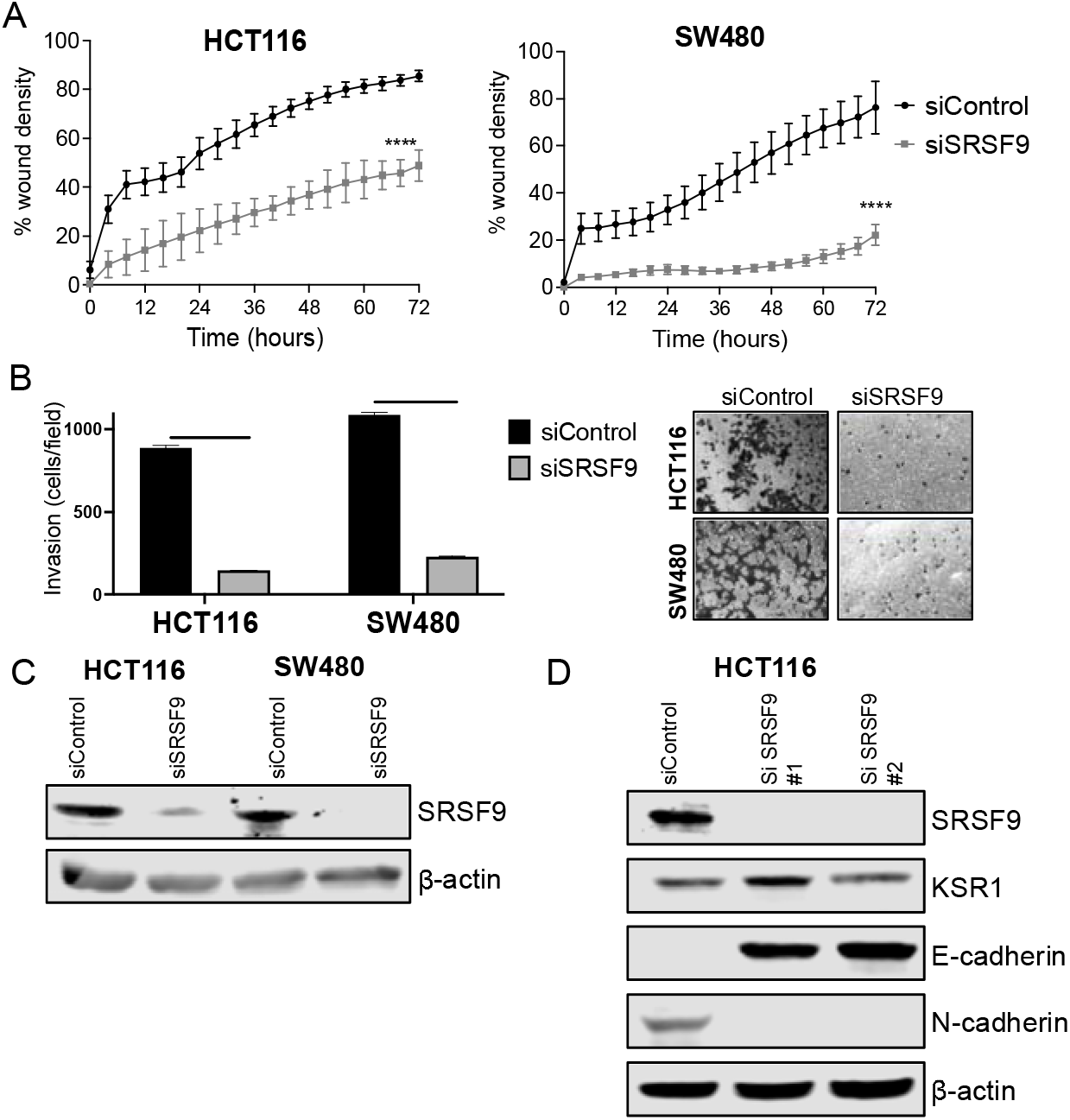
Inhibition of SRSF9 suppresses migration, invasion and mediates E-to N-cadherin switch. (A) Control or SRSF9 RNAi (siSRSF9) in HCT116 and SW480 cells were subjected to the 96-well IncuCyte scratch wound assay. The graph represents the time kinetics of percent wound density, calculated by IncuCyte ZOOM software, shown as mean ± SD, n=4; ****, P < 0.0001 (B) Control or SRSF9 RNAi (siSRSF9) in HCT116 and SW480 cells were subjected to Transwell migration assay through Matrigel^®^. The number of invaded cells per field were counted, (n=4); ****, P < 0.0001. Representative microscopic images of the respective cells following invasion through Matrigel^®^ are shown. (C) Knockdown of SRSF9 was confirmed in HCT116 and SW480 cells by Western Blot. (D) SRSF9, E-cadherin, N-cadherin, KSR1 and β actin expression following SRSF9 knockdown in HCT116 cells.

### KSR1 RNAi causes mRNA with differentially spliced exons to accumulate in polysomes

We analyzed differentially spliced events (DSEs) between KSR1 RNAi *vs*. control, in our polysome profiling RNA-seq data using rMATS 4.1.0 [35], a computational tool for differential splicing analysis that uses both exonic and intronic reads to quantify a <u>p</u>ercent <u>s</u>pliced in (PSI) value for each splicing event. We filtered out DSEs with FDR < 0.05, average minimum reads ≥ 5 and differential <u>p</u>ercent <u>s</u>pliced <u>i</u>n (ΔPSI) ≥ 10% and ΔPSI ≤ 10%. Although our sequencing depth was not optimal for a comprehensive splicing analysis, we identified over 1000 DSEs in each that are alternatively spliced in HCT116 and HCT15 cells (**Figure 4A**). Almost half of these DSEs correspond to skipped exon (SE) events, followed by alternative 5’ splice sites (A5SS), retained introns (RI), alternative 3’SS (A3SS), and mutually exclusive exons (MXEs). Comparison of DSEs between total and polysomal mRNA demonstrated that KSR1 depletion enhanced the accumulation of transcripts with included and skipped exons in polysomes in both the cell lines (**Figure 4A**). EPSTI1 has four different transcript variants because of splicing of alternative exons (either exon 12 and exon 13 and/or exon 8) yielding different products. It is likely that KSR1 may be regulating the balance between specific EPSTI1 splicing isoform, that one of these isoforms may be selectively translated to enhance motility, invasion, and EMT. Because of inadequate sequencing depth, EPSTI1 was not detected as one of the significantly spliced genes in the rMATS analysis. However, we observed a difference in the splicing pattern of EPSTI1 pre-mRNA between the control and KSR1 knockdown in HCT116 cells which suggested a exon skipping events of exon 8 in the KSR1 knockdown and inclusion of exon 12 inclusion in KSR1 knockdown cells compare to control. These results indicate a likelihood that pre-mRNA splicing playing a role its increase loading on to the polysomes in a KSR1-dependent manner.

**Figure 4.**
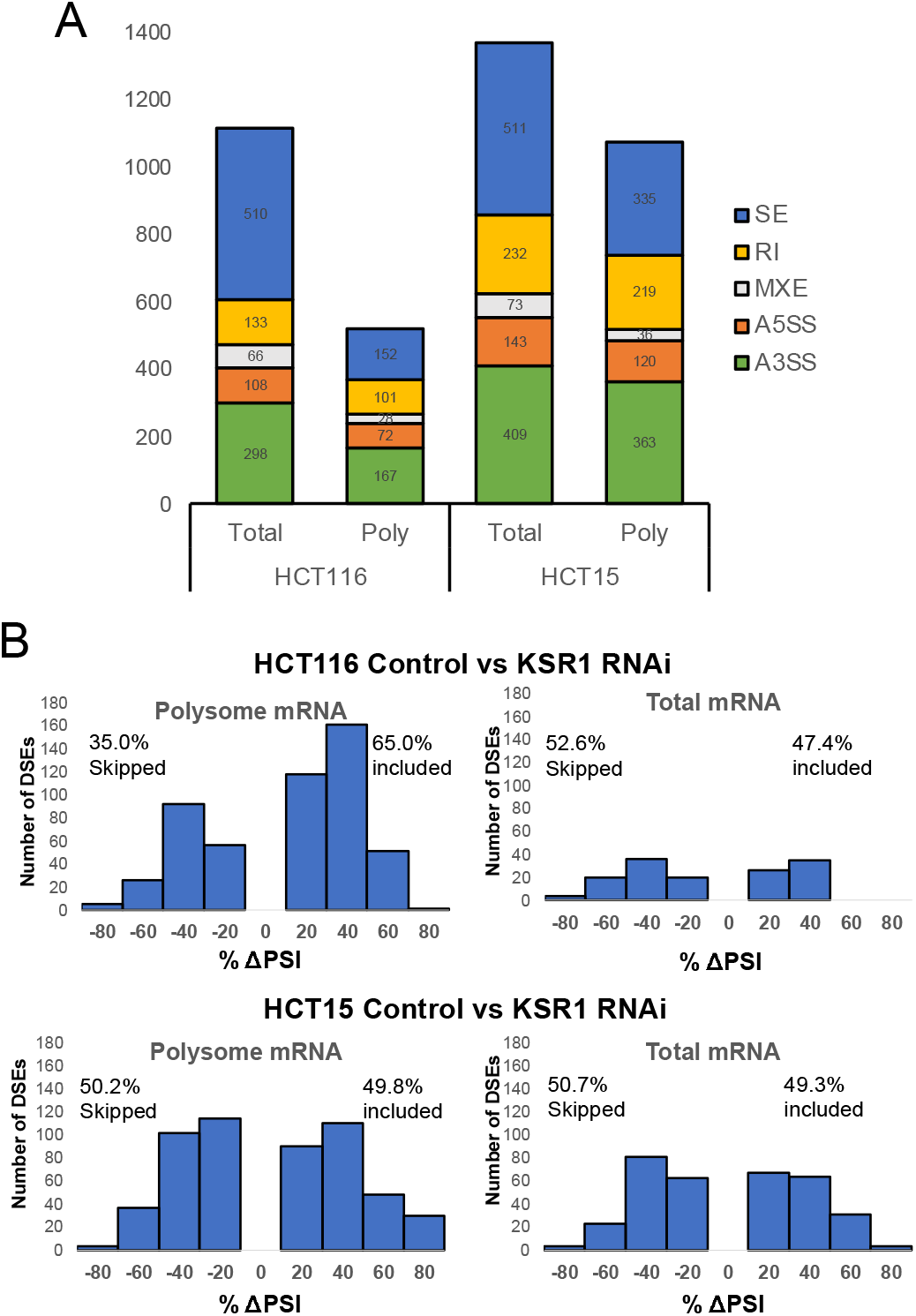
KSR1 RNAi causes mRNA with differentially spliced exons to accumulate in polysomes. (A) DSEs detected by RNA-seq in HCT116 and HCT15 KSR1 knockdown cells (n = 3; ΔPSI ≥ 10%, false discovery rate [FDR] < 5%, p < 0.01), sorted by AS event types. SE, skipped exons; RI, retained intron; MXE, mutually exclusive exon; RI, retained intron; A5SS, alternative 5’SS; A3SS, alternative 3’SS. (B) Skipped (ΔPSI %< 10%) and included (ΔPSI > 10%) exons in polysome-associated or total mRNA from KSR1 RNAi versus control HCT116 (top) and HCT15 (bottom) cells are plotted by ΔPSI values analyzed using rMATS 4.0.

### SRSF9 associates with EPSTI1 RNA

To test whether SRSF9 RNA-binding motifs reside in and around exon 8 and exon 12 of EPSTI1 RNA, we used RBPmap [36], a computational tool for mapping binding sites of RNA-binding proteins, based on *in vitro* and *in vivo* experimentally derived binding motifs. Analysis using RBPmap and visualization using the UCSC genome browser identified putative SRSF9 binding sites flanking both exons (chr13:42886388-42992249:-) (**Figure 5A**). To determine whether SRSF9 interacts with EPSTI1 mRNA, we carried out Ribonucleoprotein Immunoprecipitation (RIP) [37] experiments, which investigate changes in the association of RNAs with an RBP by immunoprecipitation analysis of native ribonucleoprotein (RNP) complexes. RIP analysis detected EPSTI1 mRNA in immunoprecipitates of SRSF9 in both HCT116 and SW480 cells (**Figure 5B**). These data suggest that SRSF9 associates with EPSTI1 mRNA and may play a role is orchestrating splicing of EPSTI1.

**Figure 5.**
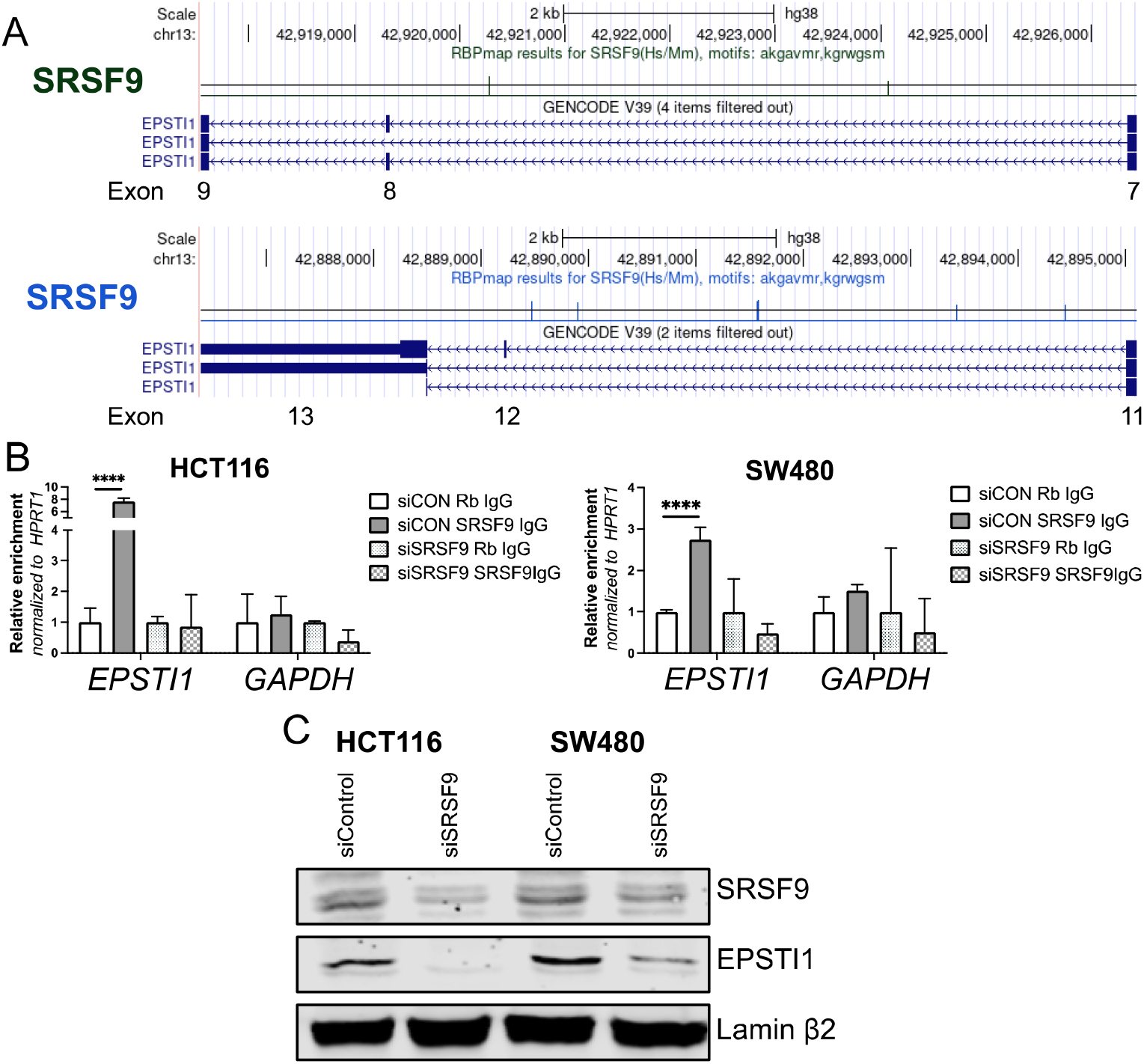
SRSF9 interacts with EPSTI1 RNA and SRSF9 inhibition suppresses EPSTI1 expression. (A) Predicted SRSF9 binding motif in flanking introns of EPSTI1 between exon 7 and exon 13 visualized using UCSC genome browser. The conserved binding motifs of SRSF9 (akgavmr, kgrwgsm) were calculated by RBPmap using high stringency and conservation filter. One colorized line represents one RBP binding motif. Longer line means higher reliability. See Table 7. (B) RNP immunoprecipitation of EPSTI1 and GAPDH mRNAs with SRSF9. The abundance of EPSTI1 and GAPDH in IPs with control IgG and anti-SRSF9 in control and SRSF9 KD HCT116 and SW480 cells. Results are normalized to HPRT mRNA and plotted as fold-enrichment of mRNA relative to IgG control samples (n=3, ****; p < 0.0001) (C) Western blot analysis of SRSF9 and EPSTI1 in control and SRSF9 knockdown (siSRSF9) in HCT116 and SW480 cells.

### Inclusion of exon 8 of EPSTI1 mRNA is optimal to CRC motility

GenBank identifies three transcript variants of EPSTI1 mRNA, a full-length transcript, an mRNA lacking exon 8, exon 12, and exon 13 (Δ*8*Δ*12*Δ*13*), and an mRNA lacking only exon 12 and exon 13 (Δ*12*Δ*13*) (**Figure 6A**). Exclusion of exon 12 creates a premature stop codon in exon 13 due to reading frame shift, producing a truncated version of the EPSTI1 (**Figure 6B**). We constructed different EPSTI1 isoforms based on the sequences deposited in GenBank using site-directed mutagenesis. We examined the ability of each variant and an artificial construct lacking only exon 8 (Δ8) to restore motility in KSR1 knockout HCT116 and SW480 cells. Measuring wound density over 72h, we found that the *EPSTI1*Δ*12*Δ*13* isoform restored migration to rates observed in control HCT116 cells (**Figure 6C**). The *EPSTI1*Δ*8*Δ*12*Δ*13* construct was approximately less than half as effective while the *EPSTI1*Δ*8* construct has no ability to rescue motility. These data suggest exon 8 inclusion optimizes the function of EPSTI1 in CRC EMT.

**Figure 6.**
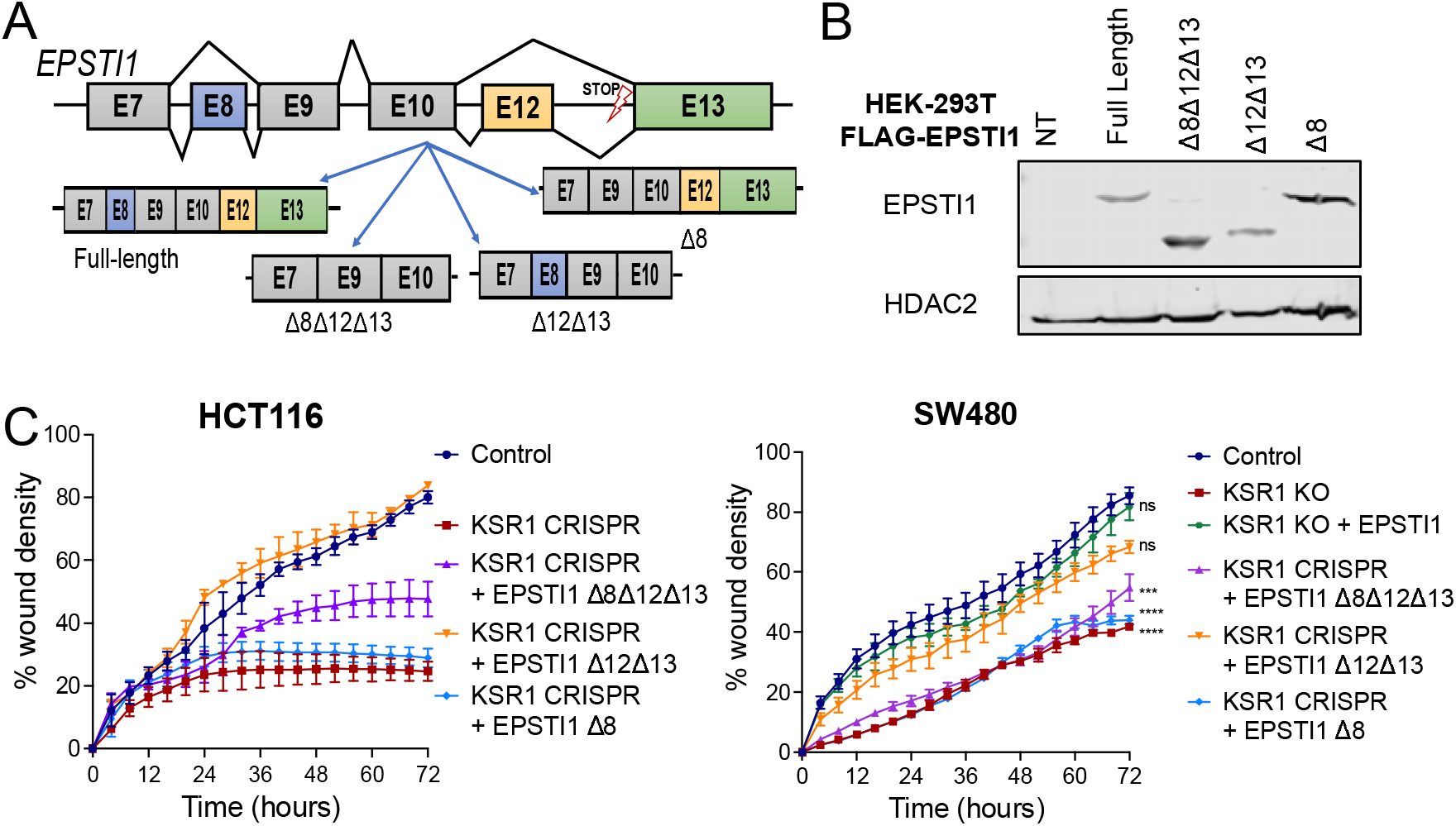
EPSTI1 exon 8 inclusion increases HCT116 cell motility. (A) Visualization of four alternative spliced forms of EPSTI1 pre-mRNA. Exons are depicted in boxes and introns in black lines. (B) Western Blot of the full-length, Δ8Δ12Δ13, Δ12Δ13, and Δ8 spliced form of EPSTI1 in HEK-293T cells (C) Control, KSR1KO, and KSR1KO cells expressing Δ8Δ12Δ13, Δ12Δ13, and Δ8 EPSTI1 HCT116 and SW480 cells were subjected to the 96-well IncuCyte scratch wound assay. The graph represents the time kinetics of percent wound density, calculated by IncuCyte ZOOM software, shown as mean ± SD, n=4.

## Discussion

Tumor cells modify the messages encoding key proteins, changing or enhancing protein function to make tumor cells more lethal. The discovery that KSR1 regulates EPSTI1 to promote the EMT-like phenotype raises new questions regarding how the translation of EPSTI1 is regulated. Post-transcriptional mechanisms and pre-mRNA splicing contributes to the plasticity required for EMT and establishment of more aggressive neoplasms [12]. Splicing factors and AS are critical points in gene expression regulation but their role in EMT is incompletely explored.

Using the computational tool for identifying oncogenic binding proteins regulating preferential mRNA translation by KSR1, we observed that binding sites for the pre-mRNA splicing factor SRSF9 were enriched in KSR1-dependent and polysome-associated mRNAs. KSR1 disruption inhibits SRSF9 expression and promotes protein degradation. SRSF9 RNAi inhibits invasion of CRC cells coincident with its ability to decrease N-cadherin expression and upregulate E-cadherin in HCT116 and SW480 cells. We further validated that SRSF9 associates with EPSTI1 mRNA, one of the KSR1-dependent targets that was identified critical for EMT-like phenotype in CRC cells and inclusion of exon 8 optimizes the function of EPSTI1 in CRC EMT-like phenotype. Our preliminary splicing analysis from the polysome profiling RNA seq data suggest that KSR1 regulates alternative splicing events, particularly exon skipping. However, the lack of sequencing depth from the RNA seq analysis does not allow us to draw detailed conclusions regarding its role in EMT-like behavior in general or EPSTI1 translation in particular. Our future directions are aimed toward determining if, and how, KSR1 regulates EPSTI1 alternative splicing and whether a preferential splice for of EPSTI1 is translated preferentially in CRC to promote the EMT-like phenotype. Collectively these data suggest that SRSF9 may be a key component of the regulatory mechanism that monitors and directs the CRC cell phenotype toward self-renewal, motility, and invasion.

Previous studies have reported SRSF9 is an oncogenic splicing factor [24, 31]. Compared to normal tissue, SRSF9 is overexpressed in a wide range of tumors including, glioblastoma, colon adenocarcinoma, and squamous cell lung carcinoma. Some reports suggested that SRSF9 promotes colony formation and growth [24], while other have reported SRSF9 promotes migratory capacity without affecting cell proliferation [11]. SRSF9 is reported to promote growth via increased β-catenin protein synthesis [24] and via regulation of apoptotic machinery [21, 23].

Determining how SRSF9-dependent induction of AS affects EMT should yield unique mechanisms underlying tumor cell metastatic behavior. SRSF9 recognizes its splicing targets via one of its two RNA recognition motifs (RRMs) that provide the binding specificity to its targets and promote exon inclusion or skipping by interacting with the splicing machinery via its C-terminal arginine/serine-rich (RS) domain. Discrete mutations in distinct patches opposite to the putative RNA binding surface disrupt RNA binding to SR domains, including SRSF9 [24]. Mutations in SWQDLKD motif (SWQDLKD to SAAALKD) impairs RNA binding [24, 31]. Future experiments will be directed toward identifying how SRSF9 expression promotes EMT in CRC cells. It is likely that mutations in either of the RRMs will show RNA binding and processing are key to the ability of SRSF9 to regulate EMT since SRSF9 interacts with EPSTI1 mRNA and phenocopies EPSTI1 action. Inclusion of exon 8 is required for optimal EPSTI1-dependent motility and invasion; however, it is not known what form of EPSTI1 is preferentially translated to prime invasive behavior.

AS of EPSTI1 may not affect the association of its mRNA onto polysomes and SRSF9 could contribute to additional mechanisms that promote their translation, via mRNA export [15, 38]. Determining the extent of SRSF9 -dependent EPSTI1 splicing and if SRSF9 preferentially directs the specific form of EPSTI1 mRNA onto polysomes will require additional studies.

While the role of KSR1 in Ras-driven tumorigenesis has been well established, the role of KSR1 in regulation of AS modulating translation of subset of mRNAs essential for EMT is novel. We demonstrate for the first time that KSR1-dependent signaling induces post-transcriptional splicing of a key target necessary for CRC cells to promote their migratory and invasive behavior. Upon disruption of SRSF9, the tumor cells reverse the mesenchymal-like behavior to an epithelial-like phenotype and EPSTI1 was identified as one of the targets orchestrating this behavior in CRC cells. It is highly possible that SRSF9 may be targeting other EMT-related mRNAs that are preferentially translated in an oncogene-driven signaling. Investigating other mRNAs effected by SRSF9 may reveal a previously underappreciated contribution of alternative splicing to cell migration, invasion, and EMT. The discovery of previously unknown mechanisms essential for EMT in cancer cells has the potential to reveal processes necessary for metastasis that may also be vulnerable to therapeutic manipulation.

## Methods

### Cell culture

Colorectal cancer cell lines HCT116, HCT15 and SW480 were acquired from American Type Culture Collection (ATCC). HCT116 and HCT15 with KSR1 knockdown or KSR1 knockout were previously generated in the lab The cells were cultured in Dulbecco’s modified Eagle’s medium (DMEM) containing high glucose with 10% fetal bovine serum (FBS) and grown at 37°C with ambient O2 and 5% CO2. Cells were routinely tested for mycoplasma. No further authentication of cell lines was performed by the authors. Quadruple mutant KAPS (KRAS^G12D^/APC^KO^/ P53^KO^/SMAD4^KO^) tumor colon organoids obtained from the Living Organoid Biobank housed by Dr. Hans Clevers and cultured as described previously [111, 112].

### Generation of EPSTI1 isoforms

The 4 isoforms of EPSTI1 were cloned into pMSCV IRES GFP. EPSTI1 transcript variant 1, accession number BC023660, protein ID AAH23660.1 was purchased from Horizon Discovery (#MHS6278202832484), To generate Flag EPSTI1 IRES GFP, full length 410 amino acid cDNA was PCR amplified (amino acids 2-410) with an N-terminal Flag tag and with flanking restriction sites EcoRI and SalI. This was inserted into the EcoRI and XhoI sites of expression vector pMSCV IRES GFP.

To generate EPST1 isoform X1 (399 amino acids) accession number XM_005266596.1, protein ID XP_005266653, amino acids 220-230 were removed (Δ exon 8) from transcript variant 1 using Quikchange site directed mutagenesis, and Flag EPSTI1 tv1 pMSCV IRES GFP as the template resulting EPSTI1 TV X1 pMSCV IRES GFP.

To generate EPSTI1 transcript variant 2 (Δ exons 8,12, and 13) accession number NM_033255.3, protein ID NP_150280, Flag EPSTI1 TV X1 pMSCV IRES GFP was used as a template to PCR amplify amino acids 2-307. The N-terminal Flag was included as well, and the EcoRI-SalI fragment was inserted into the EcoRI and XhoI sites of expression vector pMSCV IRES GFP resulting in Flag EPSTI1 TV2 pMSCV IRES GFP.

To generate EPSTI1 transcript variant 3 (Δ exons 12 and 13), accession number NM_001330543.2, protein ID NP_001317472.1, Flag EPSTI1 TV1 pMSCV IRES GFP was used as a template to PCR amplify amino acid acids 2-318 including the N-terminal Flag, and the product was inserted into the EcoRI and XhoI sites of pMSCV IRES GFP as an EcoRI-SalI fragment, resulting in Flag EPSTI TV3 pMSCV IRES GFP. All the above inserts/ deletion were Sanger sequence verified. Sequences for the PCR primers are as follows: EPSTI1-5P (5’TGTGAATTCGCCACCATGGACTACAAAGACGATGACGACAAGAACACCCGCAATAGAGTG), EPSTI1-3P (5’TCGCGGCCGCGTCGACTTAGAAAAATAATGTAGCATTTC), EPSTI TV2/3-3P (5’GAAGTCGACTCATATACCCCAGCTGTTACCGCTATTCATATTC), EPSTI1 ΔEX8-F (5’CCCACAATCCTCAACATGGGCCAGAAGCTGGG), EPSTI1 ΔEX8-R (5’CCCAGCTTCTGGCCCATGTTGAGGATTGTGGG).

All the above constructs were transfected into Phoenix GP cells using trans-lentiviral packaging system (ThermoFisher Scientific). The virus was collected, and the medium was replaced 48 hours post transfection. KSR1-CRISPR HCT116 and SW480 cells were infected with virus plus 8 μg/mL of Polybrene for 96 hours. Cells expressing EPSTI1 transgenes were selected for GFP fluorescence using fluorescence-activated cell sorting (FACS) and confirmed by Western Blotting. Sequences for PCR primers are listed below.

### RNA interference

Approximately 500,000 cells were transfected using a final concentration of 20 nM SRSF9 (Cat# 015094-09-0020, Cat# 015094-12-0020) or non-targeting (D-001810-01-20 and D-001810-02-20) ON-TARGETplus siRNAs from GE Healthcare Dharmacon using 20 μL of Lipofectamine RNAiMAX (ThermoFisher #13778-150) and 500 μL OptiMEM (ThermoFisher #31985070). Cells were incubated for 72 hours before further analysis.

### Cell lysis and western blot analysis

Whole cell lysate was extracted in radioimmunoprecipitation assay (RIPA) buffer containing 50 mM Tris-HCl, 1% NP-40, 0.5% Na deoxycholate, 0.1% Na dodecyl sulfate, 150 mM NaCl, 2 mM EDTA, 2 mM EGTA, and 1X protease and phosphatase inhibitor cocktail (Halt, ThermoFisher Scientific #78440). Cytoplasmic and nuclear fractionation was performed using NE-PER™ Nuclear and Cytoplasmic Extraction Reagents (ThermoFisher Scientific #PI78835). The estimation of protein concentration was done using BCA protein assay (Promega #PI-23222, PI-23224). Samples were diluted using 1X sample buffer (4X stock, LI-COR #928-40004) with 100 mM dithiothreitol (DTT) (10X stock, 1mM, Sigma #D9779-5G). The protein was separated using 8-12 % SDS-PAGE and transferred to nitrocellulose membrane. The membrane was blocked with Odyssey TBS blocking buffer (LICOR-Biosciences #927-50003) for 45 minutes at room temperature, then incubated with primary antibodies (*Key Resources Table*) at least overnight at 4°C. IRDye 800CW and 680RD secondary antibodies (LI-COR Biosciences # 926-32211, # 926-68072) were diluted 1:10,000 in 0.1% TBS-Tween and imaged on the Odyssey Classic Scanner (LI-COR Biosciences).

Primary antibodies were diluted as follows: SRSF9 (Cat# RN081PW, MBL BIO) 1:1000, KSR1 (Cat# ab68483, Abcam) 1:1000, EPSTI1 (Cat# 11627-1-AP, RRID: AB_2877786, Proteintech) 1:1000, N-cadherin(Cat# 13A9, Gift from Dr. Keith Johnson, UNMC) 1:20, E-cadherin(Cat# 4A2, Gift from Dr. Keith Johnson, UNMC) 1:10, Lamin β2 (Cat# A6483, RRID: AB_2767083, Abclonal) 1:2000, β actin (Cat# 47778, RRID:AB_2714189, Santa Cruz) 1:2000, phospho RSK S380 (Cat# 9341, RRID: AB_330753, Cell Signaling Technology) 1:500, Total RSK (Cat# 9355, RRID: AB_659900, Cell Signaling Technology) 1:1000.

### Splicing analysis

rMATS analysis was performed using rMATS *turbo v4.1.0* starting from the *aligned.out.bam files* obtained by mapping the reads to Homo sapiens (human) genome assembly GRCh38 (hg38) using STAR v2.7 alignment. Gencode *v22* was used for the transcript/gene annotations.

□txt files were used to pass grouping of inputs (case and control) for each condition in each cell line (HCT15/HCT116) to rMATS respectively using the parameters “single” for single-end data [-t single], read length of 76 bp [--readLength 76 bp] and allowing reads with lengths that differ from –readLength to be processed [--variable-read-length]. The detection of novel splice sites (unannotated splice sites) was enabled [--novelSS].

The final output files [AS_Event]. MATS.JC.txt and [AS_Event].MATS.JCEC.txt were combined for further analysis. For visualization of differentially spliced exons, sashimi plots were generated using rmats2sashimiplot.

#### rMATS Command

rmats.py --b1 /path/to/case.txt --b2 /path/to/control.txt --gtf /path/to/the.gtf -t single --readLength 76 --variable-read-length --nthread 15 --tstat 15 --novelSS --od /path/to/output --tmp /path/to/tmp_output

### PRADA

To predict the RNA binding proteins (RBPs) modulated upon KSR1 knockdown, a computational framework called Prioritization of Regulatory Pathways based on Analysis of RNA Dynamics Alternations (PRADA) was applied as previously described [29]. PRADA is a customized variation on lasso regression (least absolute shrinkage and selection operator) that predicts RBPs whose differential expression explains changes in the expression of their targets that is observed in the data. Briefly, genes with RefSeq ID starting with NM (Curated mRNA) were retained for the analysis. We input the log_2_ fold change (TE) values and p-values identified in the *Anota2seq* analysis from HCT116 and HCT15 cells and a RBP target matrix was created by scanning the RBP motif across the *.fasta* sequence of the mRNA targets of interest. The resulting binary matrix was created where the rows are transcripts, and the column are RBPs, and the presence of a putative binding site for RBP on the transcript is set to ‘1’ or ‘0’ otherwise. The penalties were defined as 1/|log_2_TE|. RBPs whose expression is informative for predicting the expression of their putative regulon were identified using *glmnet* R package that fits generalized linear via penalized maximum likelihood. The regularization path is computed for the lasso at a grid of values (on the log scale) for the regularization parameter lambda. The RBPs with the largest assigned coefficients (absolute value) are prioritized and plotted.

### TCGA and CPTAC data analysis

Data table was obtained from TCGA data by the UALCAN portal [39] and plotted using GraphPad Prism. Data were plotted using GraphPad Prism. Protein expression in individual stages of colon cancer were obtained from CPTAC. Z-values represent SD from the median across samples for the given cancer type. Log2 Spectral count ratio values from CPTAC were first normalized with each sample profile, then normalized across samples. Data table plotted using GraphPad Prism.

### RT-qPCR

Cells were harvested using 1 mL TRIzol (ThermoFisher Scientific #15596026) and RNA extraction was performed using RNeasy spin columns (Qiagen #74104). RNA was eluted with nuclease-free water. The RNA was quantified using a NanoDrop 2000 (Thermo Scientific) and Reverse Transcription (RT) was performed with 2 μg RNA per 40 μl reaction mixture using iScript Reverse Transcription Supermix (Bio-Rad #170-8891). RT-qPCR was performed using primers antibodies (Key Resources Table), and all targets were amplified using SsoAdvanced Universal SYBR green Supermix (Bio-Rad #1725271) with 40 cycles on a QuantStudioTM 3 (ThermoFisher Scientific). The analysis was performed using 2-ΔΔCT method [40].

### Ribonucleoprotein immunoprecipitation

Ribonucleoprotein immunoprecipitation (RIP) was performed as previously described [37]. Briefly, cells were lysed in 20□ mM Tris/HCl (pH 7.5) containing 100mM KCl, 5 mM MgCl2, 0.5 % NP-40, and the Halt Protease and Phosphatase Inhibitor Cocktail (ThermoFisher #78442) and RNase Inhibitors (ThermoFisher, # EO0381). We performed immunoprecipitations with precoated anti-SRSF9 or rabbit normal IgG-protein A Sepharose beads (GE Healthcare, catalog number: 17-1279-02) for 2 □ hours at 4□°C. The immunoprecipitated protein-RNA complexes were washed three times with 50 mM Tris HCl, pH 7.5, 150 mM NaCl, 1 mM MgCl2, 0.05% NP-40. Total RNAs were purified from the washed beads using Trizol (Qiagen) and subjected to RT-qPCR analysis for quantification.

### Cell migration (scratch-test) assay

An *in vitro* scratch test was performed with the IncuCyte Zoom according to the manufacturer’s instructions. Approximately 35,000 cells were seeded onto a 96-well ImageLock plates (Essen BioScience #4379) and grown to 90-95% confluency. The scratches were created using WoundMaker (Essen BioScience #4563) in all the wells, after which the cells were washed with 1x PBS, and media without containing serum was replaced. Images of the cells were obtained every 20 minutes for a total duration of 72 hours using IncuCyte Kinetic Live Cell Imaging System (Essen BioScience) and analyzed using the IncuCyte Zoom software (Essen BioScience). IncuCyte Software was used to calculate the relative wound density metric to quantify the cell migration over time. The metric is designed to be zero at t=0 and 100% when cell density inside the wound is the same as the cell density outside the initial wound, thus, allowing to experimentally quantify the effects of cell migration separate from changes that occurs as result of cell proliferation.

### Cell invasion (Transwell) assay

Transwell inserts (24-well Millicell cell culture, #MCEP24H48) were coated with 50 μl of Matrigel®(Corning, # 356234) and allowed to solidify for 15-30 minutes. Approximately 20,000 stably generated knockout cells, or cells after 48 hours of transfection were plated in serum free media in the upper chamber of transwell insert. Cells were allowed to invade toward 10% serum containing media in the lower chamber for 24 hours, after which cells and gel in the upper chamber was gently removed with a sterile cotton applicator and the cells in the lower side of the insert was fixed with 3.7% formaldehyde for two minutes, permeabilized with 100% methanol for 20 minutes and stained with Giemsa for 15 minutes. The numbers of cells were counted using an inverted microscope at x20 magnification.

## References

1. Gebauer, F., et al., RNA-binding proteins in human genetic disease. Nat Rev Genet, 2021. 22(3): p. 185–198.

2. Wang, E.T., et al., Alternative isoform regulation in human tissue transcriptomes. Nature, 2008. 456(7221): p. 470–6.

3. Wahl, M.C., C.L. Will, and R. Luhrmann, The spliceosome: design principles of a dynamic RNP machine. Cell, 2009. 136(4): p. 701–18.

4. Hattori, A., K. Buac, and T. Ito, Regulation of Stem Cell Self-Renewal and Oncogenesis by RNA-Binding Proteins. Adv Exp Med Biol, 2016. 907: p. 153–88.

5. Kalsotra, A. and T.A. Cooper, Functional consequences of developmentally regulated alternative splicing. Nature reviews. Genetics, 2011. 12(10): p. 715–729.

6. Chabot, B. and L. Shkreta, Defective control of pre-messenger RNA splicing in human disease. J Cell Biol, 2016. 212(1): p. 13–27.

7. Urbanski, L.M., N. Leclair, and O. Anczuków, Alternative-splicing defects in cancer: Splicing regulators and their downstream targets, guiding the way to novel cancer therapeutics. Wiley Interdiscip Rev RNA, 2018. 9(4): p. e1476.

8. Anczukow, O., et al., SRSF1-Regulated Alternative Splicing in Breast Cancer. Mol Cell, 2015. 60(1): p. 105–17.

9. Das, S., et al., Oncogenic splicing factor SRSF1 is a critical transcriptional target of MYC. Cell Rep, 2012. 1(2): p. 110–7.

10. Dvinge, H., et al., RNA splicing factors as oncoproteins and tumour suppressors. Nat Rev Cancer, 2016. 16(7): p. 413–30.

11. Park, S., et al., Differential Functions of Splicing Factors in Mammary Transformation and Breast Cancer Metastasis. Cell Rep, 2019. 29(9): p. 2672–2688 e7.

12. Pradella, D., et al., EMT and stemness: flexible processes tuned by alternative splicing in development and cancer progression. Mol Cancer, 2017. 16(1): p. 8.

13. Anczukow, O., et al., The splicing factor SRSF1 regulates apoptosis and proliferation to promote mammary epithelial cell transformation. Nat Struct Mol Biol, 2012. 19(2): p. 220–8.

14. Venables, J.P., et al., MBNL1 and RBFOX2 cooperate to establish a splicing programme involved in pluripotent stem cell differentiation. Nat Commun, 2013. 4: p. 2480.

15. Long, J.C. and J.F. Caceres, The SR protein family of splicing factors: Master regulators of gene expression. Biochemical Journal, 2009. 417(1): p. 15–27.

16. Bradley, T., M.E. Cook, and M. Blanchette, SR proteins control a complex network of RNA-processing events. Rna, 2015. 21(1): p. 75–92.

17. Paradis, C., et al., hnRNP I/PTB can antagonize the splicing repressor activity of SRp30c. RNA, 2007. 13(8): p. 1287–300.

18. Young, P.J., et al., SRp30c-dependent stimulation of survival motor neuron (SMN) exon 7 inclusion is facilitated by a direct interaction with hTra2 beta 1. Hum Mol Genet, 2002. 11(5): p. 577–87.

19. Wang, Y., et al., Tau exons 2 and 10, which are misregulated in neurodegenerative diseases, are partly regulated by silencers which bind a SRp30c.SRp55 complex that either recruits or antagonizes htra2beta1. J Biol Chem, 2005. 280(14): p. 14230–9.

20. Simard, M.J. and B. Chabot, SRp30c is a repressor of 3’splice site utilization. Mol Cell Biol, 2002. 22(12): p. 4001–10.

21. Ha, J., et al., SRSF9 Regulates Cassette Exon Splicing of Caspase-2 by Interacting with Its Downstream Exon. Cells, 2021. 10(3).

22. Oh, J., et al., Opposite Roles of Tra2beta and SRSF9 in the v10 Exon Splicing of CD44. Cancers (Basel), 2020. 12(11).

23. Cloutier, P., et al., Antagonistic effects of the SRp30c protein and cryptic 5’ splice sites on the alternative splicing of the apoptotic regulator Bcl-x. J Biol Chem, 2008. 283(31): p. 21315–24.

24. Fu, Y., et al., SRSF1 and SRSF9 RNA binding proteins promote Wnt signalling-mediated tumorigenesis by enhancing beta-catenin biosynthesis. EMBO Mol Med, 2013. 5(5): p. 737–50.

25. Rao, C., et al., KSR1-and ERK-dependent translational regulation of the epithelial-to-mesenchymal transition. Elife, 2021. 10.

26. Loh, C.Y., et al., The E-Cadherin and N-Cadherin Switch in Epithelial-to-Mesenchymal Transition: Signaling, Therapeutic Implications, and Challenges. Cells, 2019. 8(10).

27. Aiello, N.M. and Y. Kang, Context-dependent EMT programs in cancer metastasis. J Exp Med, 2019. 216(5): p. 1016–1026.

28. McCall, J.L., et al., KSR1 and EPHB4 Regulate Myc and PGC1β To Promote Survival of Human Colon Tumors. Molecular and Cellular Biology, 2016. 36(17): p. 2246–2261.

29. Yu, J., et al., RBMS1 Suppresses Colon Cancer Metastasis through Targeted Stabilization of Its mRNA Regulon. Cancer Discov, 2020. 10(9): p. 1410–1423.

30. Morris, E.J., et al., Discovery of a novel ERK inhibitor with activity in models of acquired resistance to BRAF and MEK inhibitors. Cancer Discovery, 2013. 3(7): p. 742–750.

31. Matsumoto, E., et al., AMP-activated protein kinase regulates beta-catenin protein synthesis by phosphorylating serine/arginine-rich splicing factor 9. Biochem Biophys Res Commun, 2021. 534: p. 347–352.

32. Ninomiya, K., et al., m(6) A modification of HSA TIII lncRNAs regulates temperature-dependent splicing. EMBO J, 2021. 40(15): p. e107976.

33. Akimov, V., et al., UbiSite approach for comprehensive mapping of lysine and N-terminal ubiquitination sites. Nat Struct Mol Biol, 2018. 25(7): p. 631–640.

34. Hornbeck, P.V., et al., PhosphoSitePlus, 2014: mutations, PTMs and recalibrations. Nucleic Acids Res, 2015. 43(Database issue): p. D512–20.

35. Shen, S., et al., rMATS: robust and flexible detection of differential alternative splicing from replicate RNA-Seq data. Proc Natl Acad Sci U S A, 2014. 111(51): p. E5593–601.

36. Paz, I., et al., RBPmap: a web server for mapping binding sites of RNA-binding proteins. Nucleic Acids Res, 2014. 42(Web Server issue): p. W361–7.

37. Martindale, J.L., M. Gorospe, and M.L. Idda, Ribonucleoprotein Immunoprecipitation (RIP) Analysis. Bio Protoc, 2020. 10(2): p. e3488.

38. Twyffels, L., C. Gueydan, and V. Kruys, Shuttling SR proteins: more than splicing factors. FEBS J, 2011. 278(18): p. 3246–55.

39. Chandrashekar, D.S., et al., UALCAN: A Portal for Facilitating Tumor Subgroup Gene Expression and Survival Analyses. Neoplasia, 2017. 19(8): p. 649–658.

40. Schmittgen, T.D. and K.J. Livak, Analyzing real-time PCR data by the comparative C(T) method. Nat Protoc, 2008. 3(6): p. 1101–8.

